# The Maintenance of Deleterious Variation in Wild Chinese Rhesus Macaques

**DOI:** 10.1101/2023.10.04.560901

**Authors:** Camille Steux, Zachary A. Szpiech

## Abstract

Understanding how deleterious variation is shaped and maintained in natural populations is important in conservation and evolutionary biology, as decreased fitness caused by these deleterious mutations can potentially lead to an increase in extinction risk. It is known that demographic processes can influence these patterns. For example, population bottlenecks and inbreeding increase the probability of inheriting identical-by-descent haplotypes from a recent common ancestor, creating long tracts of homozygous genotypes called runs of homozygosity (ROH), which have been associated with an accumulation of mildly deleterious homozygotes. Counter intuitively, positive selection can also maintain deleterious variants in a population through genetic hitchhiking.

Here we analyze the whole genomes of 79 wild Chinese rhesus macaques across five sub-species and characterize patterns of deleterious variation with respect to ROH and signals of recent positive selection. We show that the fraction of homozygotes occurring in long ROH is significantly higher for deleterious homozygotes than tolerated ones, whereas this trend is not observed for short and medium ROH. This confirms that inbreeding, by generating these long tracts of homozygosity, is the main driver of the high burden of homozygous deleterious alleles in wild macaque populations. Furthermore, we show evidence that homozygous LOF variants are being purged. Next, we identify 7 deleterious variants at high frequency in regions putatively under selection near genes involved with olfaction and other processes.

Our results shed light on how evolutionary processes can shape the distribution of deleterious variation in wild non-human primates.

**Significance:** Our results offer insights into the relationship between demographic and evolutionary processes and the maintenance of deleterious alleles in wild rhesus macaques. We demonstrate how inbreeding and recent positive selection can contribute to the maintenance of deleterious variants in wild non-human primate populations. Given that deleterious variation can reduce individuals reproductive fitness and contribute to extinction risks, this study is particularly relevant in the context of conservation of wild endangered species.

## Introduction

In conservation genomics, investigating patterns of deleterious variation is of crucial interest for understanding the consequences of these mutations on the phenotype and fitness of individuals, and thus on populations health and resilience (Robinson et al. 2023; Agrawal and Whitlock 2012; Bertorelle et al. 2022). It is known that the presence and accumulation of deleterious mutations in the genome, genetic load, can lead to a decrease in individual fitness (Muller 1950; Bertorelle et al. 2022). Demographic processes, such as small population size, bottlenecks, drift, selection and inbreeding can influence genetic load. For instance, small populations are more sensitive to drift which can bring mildly deleterious mutations to high frequency and even to fixation (Ohta 1973), as shown by several empirical studies that found a high genetic load in historically small populations (Xue et al. 2015; Robinson et al. 2016; de Manuel et al. 2020; Von Seth et al. 2021). Recent inbreeding can rapidly decrease the fitness of individuals by increasing the homozygosy of recessive deleterious variants which then become expressed (Hedrick and Kalinowski 2000; Charlesworth and Willis 2009; Räikkönen et al. 2009; Hedrick et al. 2014; Robinson et al. 2019; Stoffel et al. 2021; Swinford et al. 2023). On the other hand, several studies have suggested that, in the long run, inbreeding could also decrease the genetic load by purging strongly deleterious (e.g. loss-of-function, LOF) alleles from populations (Xue et al. 2015; Benazzo et al. 2017; Kleinman-Ruiz et al. 2022). As for natural selection, slightly deleterious alleles can increase in frequency by genetic hitchhiking if they are linked to positively selected loci (Charlesworth and Jensen 2021; Van Hooft et al. 2021; Barton 2000). As genetic load can potentially lead to an increase in extinction risk (Robinson et al. 2023; Lynch et al. 1995), it poses significant challenges for conservation management, particularly in small and isolated endangered populations (Robinson et al. 2023; Leon-Apodaca et al. 2023). Therefore, there is a need to better understand how evolutionary forces shape the distribution of deleterious mutations in wild populations to inform conservation efforts.

A well documented process influencing the maintenance of genetic load is the gathering of recessive deleterious variation in homozygous state via inbreeding (Szpiech et al. 2013; Zhang et al. 2015; Charlesworth and Willis 2009; Szpiech et al. 2019). Small population size and isolation promotes mating between close relatives, which generates long tracts of identical-by-descent segments in the offspring due to the presence of shared haplotypes in the parents inherited from the same recent ancestor (Woods et al. 2006; Ceballos et al. 2018). These long tracts of homozygous haplotypes are called runs of homozygosity (ROH) and, given the relationship between ROH and autozygosity (McQuillan et al. 2008; Kardos et al. 2015), ROH are useful for analyzing the effects of inbreeding in populations (Keller et al. 2011).

Recent inbreeding generates long tracts of homozygosity. Over time, mutations and recombination events break up the underlying haplotypes which can eventually result in shorter ROH (Pemberton et al. 2012; Ceballos et al. 2018). Therefore, the length of ROH in an individual can be used as a proxy to infer the time of the events that generated them (Ceballos et al. 2018), and the study of these regions can also be informative about long-term population demographic history (Pemberton et al. 2012; Ceballos et al. 2018; Pemberton and Szpiech 2018).

It is known that increased homozygosity can lead to a decrease in an offspring’s fitness, a phenomenon called inbreeding depression (Woods et al. 2006; Darwin 1877). Studies in humans have shown that long ROH are a reservoir of homozygous deleterious variation, meaning that an individual with a high burden in ROH is likely to have a high burden in deleterious homozygotes as well (Szpiech et al. 2013; Pemberton and Szpiech 2018; Szpiech et al. 2019). In dogs, ROH have been associated with some medical conditions in specific breeds, such as lymphoma in Labradors and Golden Retrievers (Mooney et al. 2021). Indeed several studies have shown a significant association between Wright’s inbreeding coefficient and some complex traits (Rudan et al. 2003). In natural populations, despite that some endangered populations challenge this paradigm (Robinson et al. 2016; Benazzo et al. 2017; Robinson et al. 2019; Leon-Apodaca et al. 2023), several studies have shown that inbreeding reduces individuals reproduction and survival, and that inbreeding depression can increase species extinction risk (Brüniche-Olsen et al. 2018; Spielman et al. 2004; Räikkönen et al. 2009; Hedrick et al. 2014; Robinson et al. 2019).

Here, we assess the accumulation and distribution of deleterious and LOF variation found in a sample of wild non-human primates in the context of demographic and selective forces. Liu et al. (2018) have sequenced the whole genome of 79 wild Chinese rhesus macaques covering the five currently recognised subspecies (*M. m. brevicaudus, M. m. lasiotis, M. m. littoralis, M. m. mulatta*, and *M. m. tcheliensis*), and investigated their past evolutionary history. After diverging from Indian rhesus macaques around 162 kya and colonizing the South of China (Hernandez et al. 2007), Liu et al. (2018) inferred a subsequent step-by-step divergence scenario for Chinese rhesus macaques, with the subspecies successively diverging from the ancestral population while passing through population bottlenecks. *M. m. mulatta* diverged first from the ancestral population around 126 kya, followed by *M. m. lasiotis* 104 kya, then *M. m. brevicaudus* 62 kya, and finally *M. m. littoralis* and *M. m. tcheliensis* diverging from each other 51 kya. Furthermore, Liu et al. (2018) found signals of morphological and physiological adaptation to habitat among the subspecies, and found several human disease-causing variants, some of them being private to some subspecies. While Liu et al. (2018) discussed at length the implications of their results for human medical research, here we explore the evolutionary mechanisms that might lead to the maintenance of deleterious variation in these natural populations of rhesus macaques in a conservation context. Given that they differ in their recent demographic history, we expect to observe different genomic patterns across the subspecies, which makes this data set an opportunity to explore and compare patterns of deleterious variation in a wild non-human primate.

We investigate patterns of deleterious (and strongly deleterious) variation in relation to ROH and signals of recent positive selection. In the first part, we follow Szpiech et al. (2013) and compute the fraction of deleterious, LOF (e.g. assumed to be strongly deleterious) and tolerated variants falling within different size class ROH (short, medium and long). Then we identify regions putatively under positive selection in each of the five subspecies and identify within these regions deleterious variants at high frequency and the genes they are in. We show that both inbreeding and natural selection participate in the maintenance of deleterious variation in the genomes of the studied populations, and we also find signals of purging of LOF mutations.

## Results

### Genomic Data Set

Data was retrieved from Liu et al. (2018), who sequenced whole genomes of 79 wild Chinese rhesus macaques across the five subspecies. After lifting over to a more recent reference genome (Mmul_10, Warren et al. (2020)) and filtering and phasing (see Materials and Methods), our data set consisted in 5,713,999 single-nucleotide polymorphisms for which at least one of the 79 individuals had a non-reference allele. Variant harmfulness was predicted with SIFT 4G (Vaser et al. 2016). In total, SIFT annotated 17,477 variants with confidence across our 79 individuals (see fig. 1). Among these, 14,913 variants were predicted to be tolerated (8,728 synonymous, 6,185 non-synonymous), 2,385 to be deleterious (33 synonymous, 2,352 non-synonymous), and 165 variants were annotated as stop-gain, 14 as stop-loss and none as start-loss. We labeled stop-loss and stop-gain variants as LOF (179 sites), given that we expect such variants to likely affect transcription (see Material and Methods). LOF variants are expected to be strongly deleterious, in particular more deleterious than the variants predicted to be deleterious by SIFT (Lek et al. 2016). From now on and throughout the rest of the paper, the terms deleterious and LOF refer to the SIFT classification.

**Figure 1:**
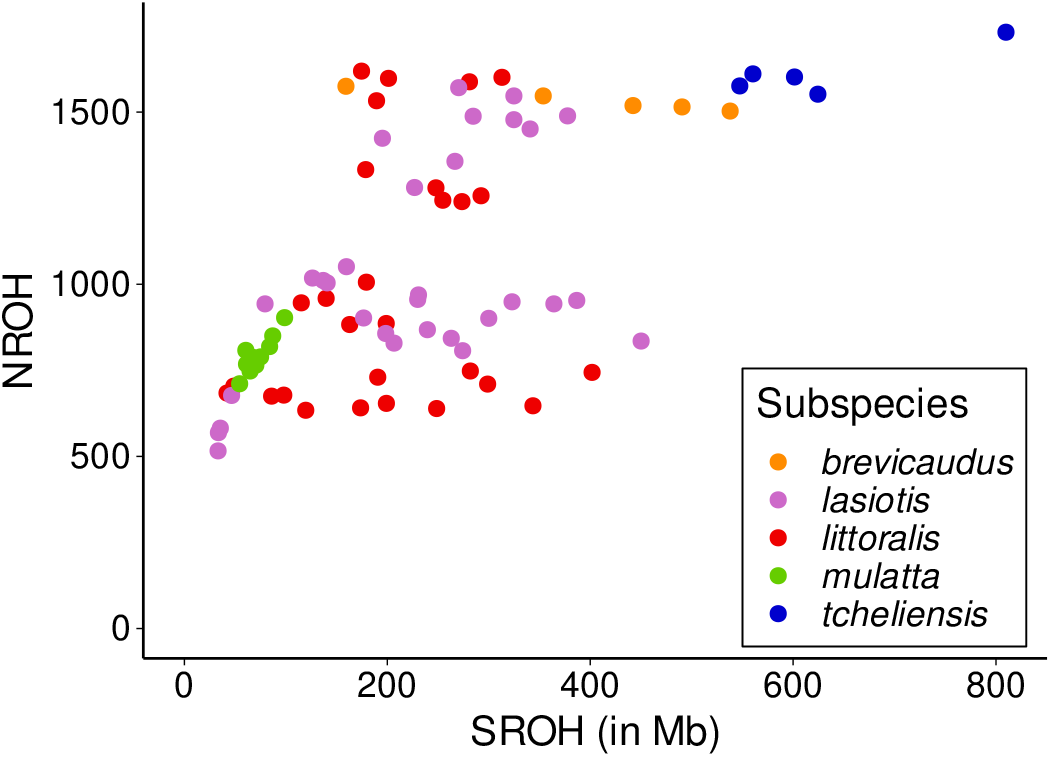
Total number of ROH (NROH) versus the total length of ROH (SROH) for each individual, colored by subspecies.

### Runs of Homozygosity

ROH were identified using GARLIC v1.1.6a (Szpiech et al. 2017). Using a gaussian mixture model, GARLIC automatically classified all ROH into three different size classes: short ROH (< 92,729 bp), medium ROH (>= 92,729 bp and < 441,568 bp) and long ROH (>= 441,568 bp). The mean length was 44,695 bp, 157,206 bp and 2.06 Mbp for short, medium and long ROH respectively. The longest identified ROH across all individuals was 30.7 Mbp long and was found in *M. m. lasiotis* (individual C_rhe_50).

To summarize the distribution of ROH, we computed two statistics used in previous studies (Pemberton et al. 2012; Ceballos et al. 2018; Szpiech et al. 2013): the sum of the total length of ROH per individual (SROH) and the total number of ROH per individual (NROH). We observe in fig. 1 different patterns among the subspecies. Individuals from the same subspecies tend to cluster together, though there is some heterogeneity for *M. m. lasiotis* and *M. m. littoralis*. The subspecies *M. m. mulatta* exhibits short total lengths of ROH and small numbers of ROH, whereas *M. m. tcheliensis* presents the longest and most numerous ROH, followed by *M. m. brevicaudus. M. m. lasiotis* and *M. m. littoralis* present similar intermediate distributions.

For each individual, we also calculated the total fraction of the genome covered by any ROH or by ROH of a specific length category: short, medium and long. Fig. 2 shows the total fraction of the genome covered by ROH in each individual. ROH genome coverage ranges from < 5% to around 28%, and the proportion of short, medium and long ROH varies across individuals. The subspecies *M. m. tcheliensis* and *M. m. brevicaudus* exhibit the individuals with the highest ROH genome coverage, suggesting a strong bottleneck or inbreeding in these subspecies, whereas individuals from *M. m. mulatta* show the lowest ROH genome coverage likely indicating a large effective population size.

**Figure 2:**
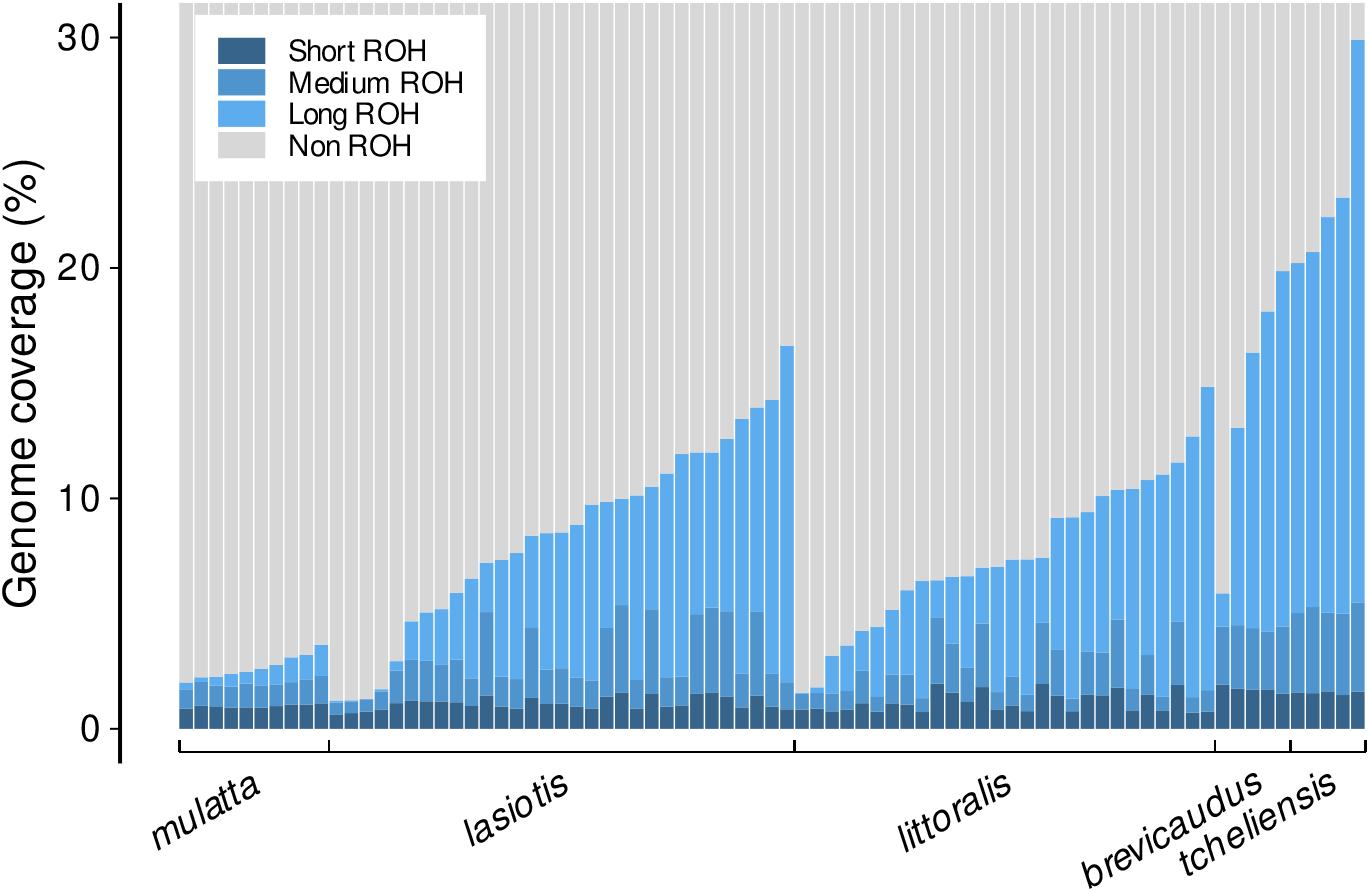
Fraction of the genome covered by ROH. Each shade of blue represents a size class ROH: short ROH in dark blue, medium ROH in medium blue and long ROH in light blue. Each bar represents one individual.

### Total distribution of deleterious variation

We used SIFT (Vaser et al. 2016) (see Material and Methods) to predict the harmfulness of variants and group them into deleterious, tolerated, and LOF categories. Fig. 3A shows that individuals carry between 23 and up to 64 deleterious alternate-allele homozygotes, with *M. m. mulatta* harboring the smallest number (compared to *M. m. lasiotis, M. m. littoralis, M. m. brevicaudus* and *M. m. tcheliensis*, Mann–Whitney–Wilcoxon, *p* = 0.0004, *p* = 0.008, *p* = 0.008 and *p* = 0.004, respectively), while the other four subspecies carry similar numbers. This pattern is consistent with the observed distribution of ROH across subspecies, as we would expect the subspecies with the lowest levels of ROH to have the lowest levels of homozygotes. Regarding deleterious heterozygotes, we observe that *M. m. tcheliensis* has the fewest number (between 160 and 215) (compared to *M. m. mulatta, M. m. lasiotis, M. m. littoralis* and *M. m. brevicaudus*, Mann–Whitney–Wilcoxon, *p* = 0.001, *p* = 0.0003, *p* = 0.0005 and *p* = 0.008, respectively), and the four other subspecies are roughly equal (between 190 and 280). Finally, the highest genetic load (2*number of homozygotes + number of heterozygotes) (fig. 3C) is found in *M. m. brevicaudus* (compared to *M. m. mulatta, M. m. lasiotis, M. m. littoralis* and *M. m. tcheliensis*, Mann–Whitney–Wilcoxon, *p* = 0.004, *p* = 0.018, *p* = 0.020 and *p* = 0.048, respectively), whereas *M. m. mulatta* carries a similar load to *M. m. tcheliensis* and a significantly smaller load compared to the other three subspecies (compared to *M. m. lasiotis, M. m. littoralis* and *M. m. brevicaudus*, Mann–Whitney–Wilcoxon, *p* = 0.003, *p* = 0.03 and *p* = 0.004, respectively). The individual with the highest number of deleterious alleles is found in *M. m. littoralis*, with 373 deleterious alleles.

**Figure 3:**
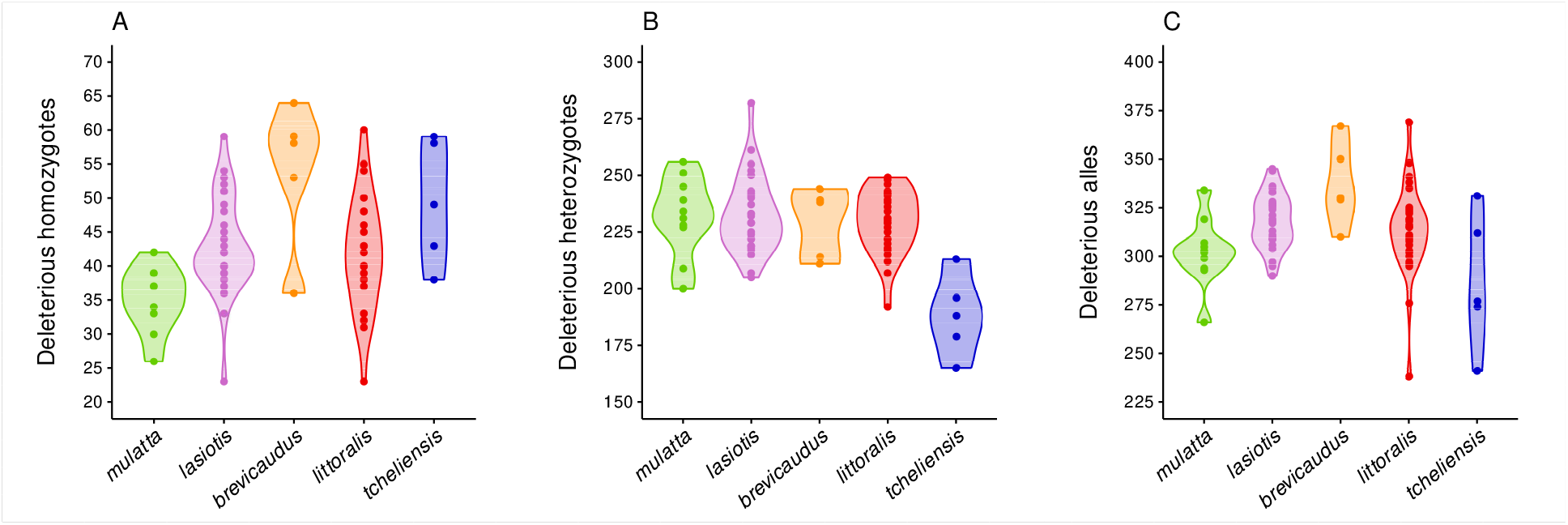
Distribution of the number of (A) Deleterious alternate-allele homozygotes, (B) Deleterious heterozygotes and (C) Deleterious alleles predicted by SIFT (Vaser et al. 2016) for each subspecies.

### Deleterious and Tolerated Homozygotes in Runs of Homozygosity

For each individual, we computed the number of deleterious alternate-allele homozygotes and tolerated alternate-allele homozygotes that occur within short, medium and long ROH. fig. 3 and Table S3 show that the number of deleterious or tolerated alternate homozygotes falling in ROH increases as we move from *M. m. mulatta* to *M. m. lasiotis, M. m. littoralis, M. m. brevicaudus* and to *M. m. tcheliensis*. This pattern across supspecies is the same as the pattern observed in ROH (see fig. 2), which is consistent with the fact that the higher the ROH genome coverage is, the higher is the probability for homozygotes to fall within these regions. Furthermore, we see that the number of both deleterious and tolerated homozygotes falling in ROH increases on average as we move from short, medium to long ROH, which is expected as ROH are defined as regions carrying more homozygotes than non ROH regions.

For each individual, we then computed the fraction of deleterious and tolerated homozygotes occurring within any size ROH and each size class ROH individually. In fig. 4A, we observe a positive correlation between both the deleterious (Pearson *r* = 0.878, *p* = 2.5 × 10^−26^) and the tolerated (Pearson *r* = 0.962, *p* = 3.9 × 10^−45^) fraction of homozygotes falling into ROH and the fraction of the genome covered by ROH. These correlations were expected, as we can expect a higher number of homozygotes falling within ROH when the length of ROH in the genome increases. Furthermore, we observe a higher fraction of deleterious homozygotes falling within ROH than the fraction of tolerated homozygotes, suggesting a faster enrichment of deleterious homozygotes per unit increase of genomic ROH coverage.

**Figure 4:**
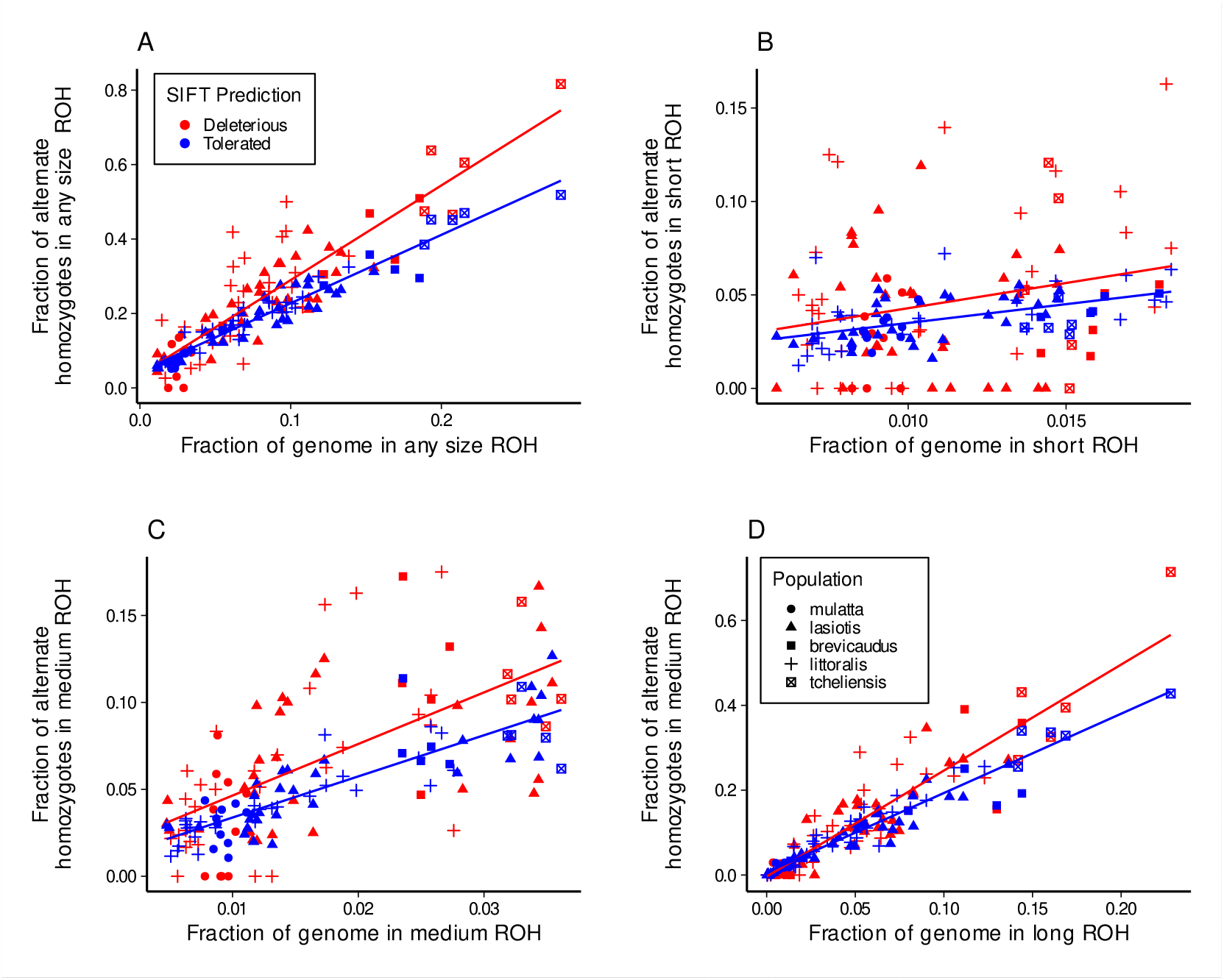
Fraction of genome-wide alternate homozygotes falling in ROH versus the fraction of the genome covered by ROH. Each point is an individual, blue points represent an individual’s deleterious homozygotes and red points represent an individual’s tolerated homozygotes. (A) Any size ROH, (B) Short ROH, (C) Medium ROH and (D) Long ROH.

To test this observation and assess the statistical significance in the difference between these trends, we fit a linear model which is described in Material and Methods (Equation 1). We found no significant difference in the intercept of the two lines (*p* = 0.61225), but a significant difference in slope (β_3_ = 0.70608, *p* = 4.9 × 10^−5^), indicating a significant higher enrichment of deleterious homozygotes per unit increase in ROH as compared to tolerated homozygotes.

Figures 4B-D represent the fraction of deleterious and tolerated homozygotes falling within short, medium, or long ROH, versus the fraction of genome covered by short, medium and long ROH, respectively. For each size class, we observe the expected and significant positive correlation between the fraction of homozygotes occurring in ROH and the fraction of the genome covered by ROH (*p* = 0.04, *p* = 6 × 10^−10^ and *p* = 1.4 × 10^−30^ for deleterious homozygotes in short, medium and long ROH, respectively, and *p* = 2.7 × 10^−07^, *p* = 3.8 × 10^−25^ and *p* = 3.3 × 10^−43^ for tolerated homozygotes in short, medium and long ROH, respectively). Furthermore, for each class and especially for long ROH, the fraction of deleterious homozygotes falling in ROH appears generally higher than the fraction of tolerated homozygotes.

We assessed the statistical significance between the two lines using the linear model described in Material and Methods (Equation 2). For short and medium ROH, there is no significant difference in intercept (*p* = 0.944 and *p* = 0.439, respectively) or slope (*p* = 0.604 and *p* = 0.181, respectively). However for long ROH, while there is no significant difference in intercept (*p* = 0.35), we found a significant difference in slope (β_3_ = 0.62870, *p* = 3.4 × 10^−5^). This indicates a significant higher enrichment of deleterious homozygotes per unit increase in long ROH compared to tolerated homozygotes. These results are consistent with the previous work of Szpiech et al. (2013), Pemberton and Szpiech (2018), and Szpiech et al. (2019), who also found a higher enrichment of deleterious homozygotes than tolerated ones in long ROH (Szpiech et al. 2013; Pemberton and Szpiech 2018; Szpiech et al. 2019).

### Loss-of-function Variants

As a critical part of this work relies on predicting whether a variant is harmful, and while deleterious and tolerated sets of variants are likely to represent, in aggregate, mutations that are deleterious or tolerated, we focus now on LOF polymorphisms that are *a priori* likely to be strongly deleterious (Lek et al. 2016). We thus compared the distribution of LOF variants (n=179) to our set of SIFT-predicted tolerated sites (n=14,913).

In fig. 5, we plot the distribution of LOF variants across the five subspecies. Remarkably, individuals carry on average between 2 and 8 LOF variants in homozygous state (fig. 5A), including one *M. m. littoralis* with 12 LOF homozygotes. This is noteworthy given that LOF variants are likely to affect protein function and thus impact individuals fitness. Individuals carry between 16 and 34 LOF in heterozygous state, *M. m. mulatta* and *M. m. lasiotis* having significantly more LOF heterozygotes than *M. m. littoralis* and *M. m. tcheliensis* (compared to *M. m. littoralis* and *M. m. tcheliensis*, Mann–Whitney–Wilcoxon, *p* = 0.015 and *p* = 0.013 for *M. m. mulatta, p* = 0.006 and *p* = 0.014 for *M. m. lasiotis*, respectively) (fig. 5B). Finally, there is no significant general trend in the genetic load (2*number of homozygotes + number of heterozygotes), though *M. m. lasiotis* has a significant higher genetic load than *M. m. littoralis* and *M. m. tcheliensis* (Mann–Whitney–Wilcoxon, *p* = 0.003 and *p* = 0.031, respectively).

**Figure 5:**
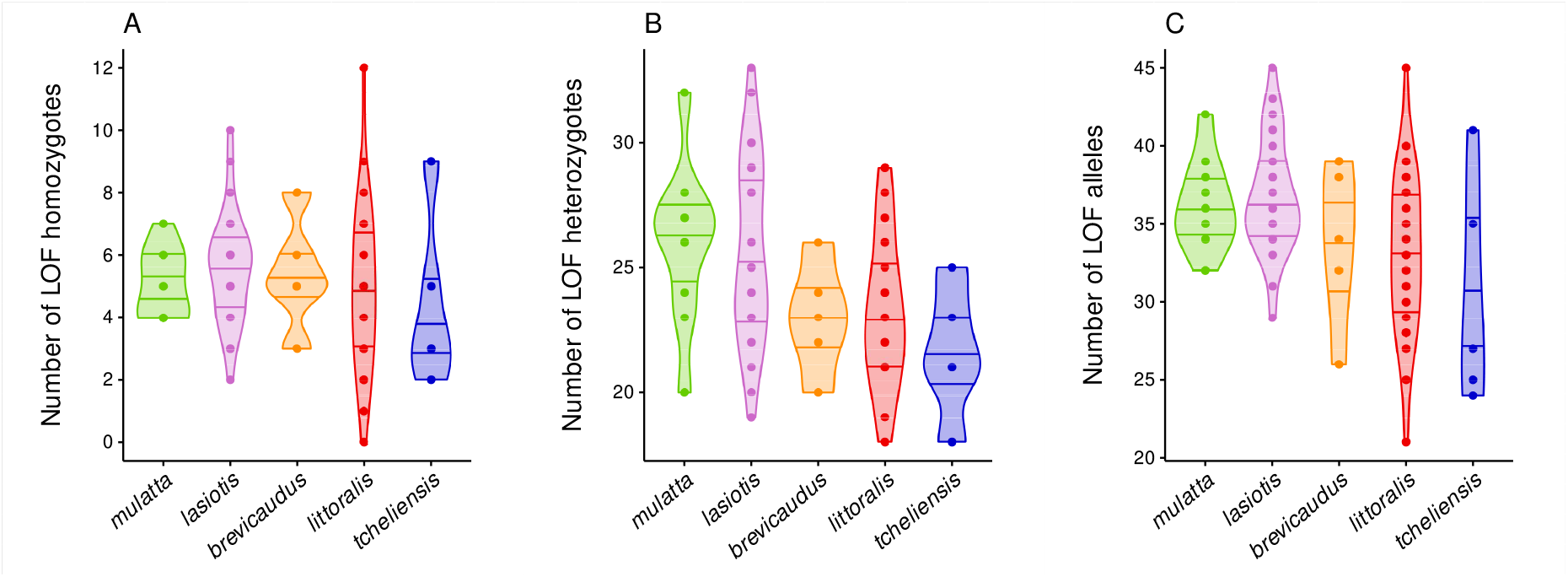
Distribution of the genome-wide number of loss-of-function (LOF) (A) Alternate homozygotes, (B) Heterozygotes and (C) Alleles predicted by SIFT (Vaser et al. 2016) for each subspecies.

For both tolerated and LOF variants, we counted the number of polymorphic sites homozygous for the alternate allele occurring inside ROH. Then, in the same way as done above, we computed, for each individual, the fraction of tolerated and LOF homozygous occurring within ROH for any size ROH and each size class individually (see Supplementary Tables S3).

Because the number of LOF variants is much smaller than the number of tolerated ones, we partitioned our individuals into two groups based on their ROH genome coverage. For any size ROH and each size class ROH, we classified individuals as Low ROH if the fraction of their genome covered by ROH was lower than the median fraction across individuals, and classified individuals as High ROH if the fraction of their genome covered by ROH was higher. We then plotted the distribution of the fraction of LOF and tolerated variants occurring in ROH for each category of individuals.

Fig. 6 shows the fraction of tolerated and LOF homozygotes falling outside or inside ROH regions, for any size ROH and each size class ROH, and for individuals classified as having a low ROH genome coverage (Low ROH) and individuals classified as having a high ROH genome coverage (High ROH). Mann–Whitney–Wilcoxon tests were performed to assess the significance of the difference between the distribution of LOF and tolerated variants.

**Figure 6:**
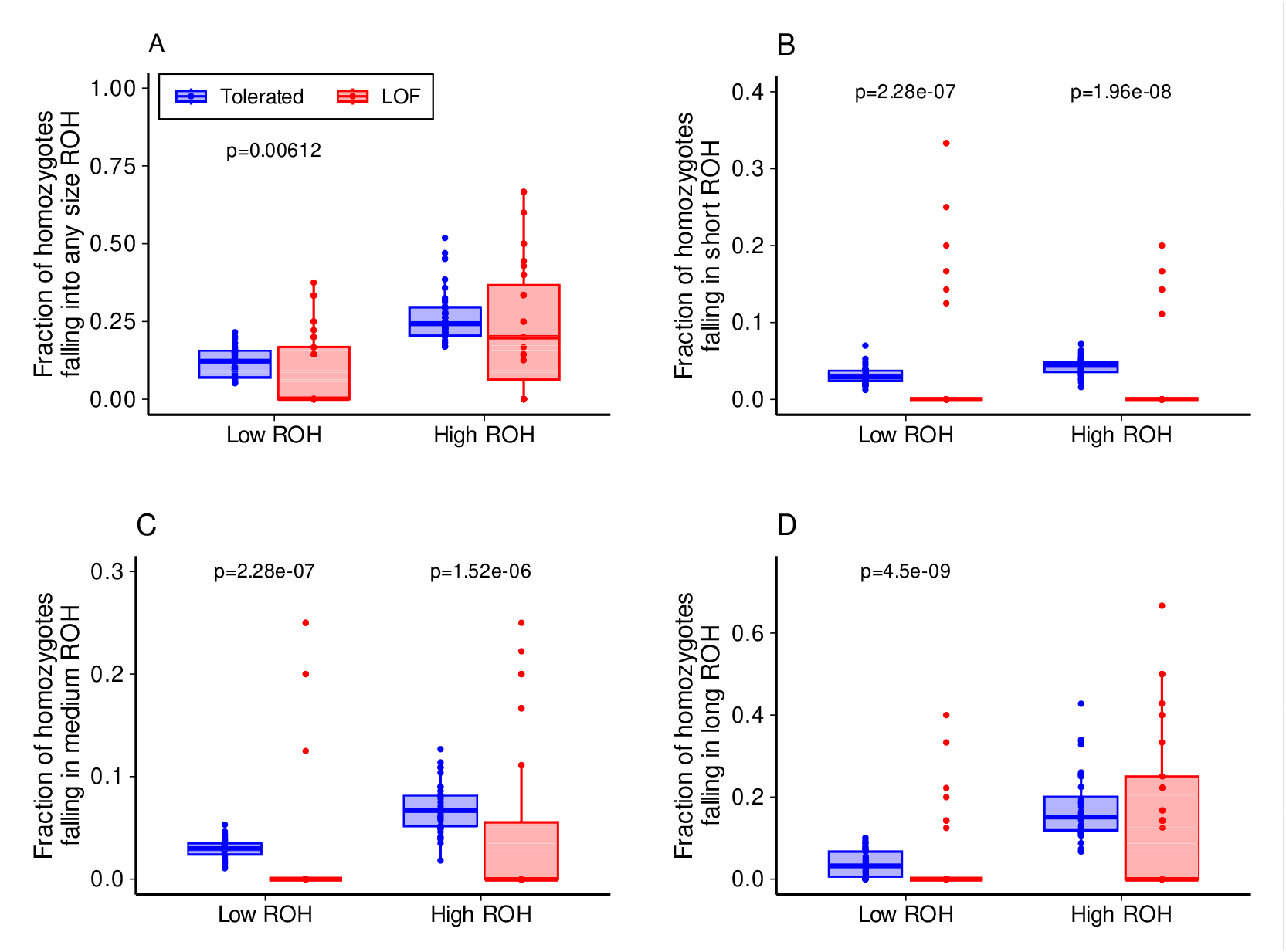
Fraction of genome-wide alternate homozygotes falling in ROH for individuals with low genomic ROH coverage (Low ROH) and individuals with a high genomic ROH coverage (High ROH). p-values correspond to Mann–Whitney–Wilcoxon tests, testing for a lower distribution of LOF variants as compared to tolerated variants. Only significant p-values are indicated. (A) Any size ROH, (B) Short ROH, (C) Medium ROH and (D) Long ROH.

For any size ROH, medium and long ROH, we observe a significantly higher fraction of homozygotes within ROH in individuals categorized as having a high ROH genome coverage than in individuals having a low ROH coverage (Mann–Whitney–Wilcoxon, *p* = 2 × 10^−12^, *p* = 0.05, *p* = 4 × 10^−12^ for any size, medium and long ROH, respectively) (fig. 6). This was expected, as more variants are predicted to fall within ROH as the total length of ROH in the genome increases. For individuals with low levels of ROH, we observe a significantly lower distribution of LOF variants compared to tolerated ones for all ROH regardless of their size (Mann–Whitney–Wilcoxon, *p* = 0.00612, *p* = 2.28 × 10^−7^, *p* = 2.28 × 10^−7^ and *p* = 4.5 × 10^−9^ for any size, short, medium and long ROH, respectively). For individuals with high levels of ROH, we observe the same significant trend for short and medium ROH (Mann–Whitney–Wilcoxon, *p* = 1.96 × 10^−8^ and *p* = 1.52 × 10^−6^ for short and medium ROH, respectively), whereas there is no significant difference in the distribution of LOF and tolerated variants in any size and long ROH (Mann–Whitney–Wilcoxon, *p* = 0.105 and *p* = 0.205 for any size and long ROH, respectively).

### Deleterious sites in regions under positive selection

We further investigated if some deleterious mutations could have been maintained by genetic hitch-hiking. We identified regions under selection by using the nSL statistic (Ferrer-Admetlla et al. 2014) as implemented in selscan (Szpiech and Hernandez 2014) in each subspecies separately. Between 237 and 254 windows of 100 kb were identified to be under selection in each subspecies (see Supplementary Tables S5-9). We then partitioned deleterious variants in two subsets, the ones falling outside regions under selection and the ones falling inside, and computed the site frequency spectrum (SFS) on each of these subsets, to test for an excess of high-frequency deleterious sites (see Supplementary fig. 4). We identified 7 deleterious mutations at high frequency (>0.7) in regions under selection across the 5 subspecies. Their positions, the genes involved, and their associated functions are summarized in Table 1.

**Table 1:** High frequency (> 0.7) deleterious sites in genomic regions identified to be under selection and the gene ontologies involved.

| Site position | Subspecies | Gene | PANTHER Family/ Sub-family | PANTHER Protein Class |
| --- | --- | --- | --- | --- |
| chr1:1218571 | *mulatta* | ENSMMUG-00000056211<br>OR11L1 | NA<br>Olfactory receptor (PTHR24242:SF386) | NA<br>G-protein coupled receptor (PC00021) |
| chr8:55470846 | *tcheliensis* | RP1 | Oxygen-regulated 1 (PTHR23005:SF4) | NA |
| chr14:59880975 | *mulatta, lasiotis, littoralis* | OR56A1 | Olfactory receptor 56A1 (PTHR26450:SF425) | Transmembrane signal receptor(PC00197) |
| chr14:95238067<br>chr14:95248391 | *brevicaudus* | CEP126 | Centrosomal protein of 126 KDA (PTHR31191:SF4) | DNA metabolism protein (PC00009) |
| chr14:95618606 | *brevicaudus* | BIRC2 | Baculoviral IAP repeat-containing 2 (PTHR10044:SF79) | Protease inhibitor (PC00191) |
| chr14:96147958 | *brevicaudus* | MMP12 | Macrophage metalloelastase (PTHR10201:SF267) | Metalloprotease (PC00153) |

We found that both *M. m. mulatta* and *M. m. tcheliensis* harbour one private mutation at high frequency in selected regions, on chromosome 1 and 8 respectively, involving an olfactory receptor (OR11L1) in the case of *M. m. mulatta* and a gene involed in microtubule binding (RP1) for *M. m. tcheliensis*. The three subspecies *M. m. mulatta, M. m. lasiotis* and *M. m. littoralis* share one high-frequency variant in selected regions on chromosome 14 at position 59,880,975, involved with the gene OR56A1 that encodes an olfactory receptor. Interestingly, we also found four variants at high-frequency in *M. m. brevicaudus* on chromosome 14 in a window of less than 1Mb, involved with the genes CEP126 (coding a DNA metabolism protein), BIRC2 (a protease inhibitor), and MMPI2 (a metalloprotease).

## Discussion

### Patterns of ROH

In the 79 individuals of our data set, we identified patterns of ROH and showed that the subspecies carry different burden of ROH. The observed patterns of ROH are consistent with what is known of the evolutionary history of Chinese rhesus macaques. Indeed, under the demographic scenario inferred by Liu et al. (2018) and presented in the Introduction, *M. m. mulatta* would have gone through only one bottleneck, whereas *M. m. tcheliensis* would have gone through four bottlenecks. As episodes of small population size promote homozygosity, this is consistent with the significantly higher burden in ROH, especially short and medium size ROH, in *M. m. tcheliensis* compared to *M. m. mulatta*. The highest total length of long ROH in *M. m. tcheliensis* is likely the result of recent small population size, which is consistent with low genetic diversity in this subspecies, compared to the other subspecies, especially *M. m. mulatta* which harbours the highest genetic diversity (Liu et al. 2018). Regarding the other three subspecies, given the demographic history inferred by Liu et al. (2018), we would have expected to observe shorter and fewer ROH for *M. m. lasiotis*, followed by *M. m. brevicaudus* and then *M. m. littoralis*, as they went through an increasing number of bottlenecks. However, these three subspecies have roughly similar levels of ROH (fig. 2), although they are intermediate to *M. m. mulatta* and *M. m. tcheliensis*, consistent with similar patterns in humans (Pemberton et al. 2012).

### Long ROH are enriched in deleterious homozygotes

In accordance with previous work (Szpiech et al. 2013, 2019; Pemberton and Szpiech 2018; Woods et al. 2006; Sams and Boyko 2019; Zhang et al. 2015), our results show that long ROH are enriched in deleterious homozygote variation. Intuitively, when the fraction of a genome covered by ROH increases, the genome gets enriched in polymorphic sites in homozygous state. We have demonstrated here that this enrichment is higher for deleterious alleles than tolerated ones in long ROH. Because deleterious variants tend to be present in low frequency in a population, it is more likely for two deleterious alleles to get together in homozygous form by being inherited together from a recent common ancestor than by chance (Szpiech et al. 2013; Pemberton and Szpiech 2018; Szpiech et al. 2019). This is also true for tolerated alleles, but to a lesser extent given that they are likely to be present in a population at a higher frequency (Gillespie 2004).

As long ROH are likely to result from mating between two closely related individuals, these results suggest that recent inbreeding is the process driving the enrichment in deleterious homozygotes in inbred populations. These observations corroborate what has been known for a long time as inbreeding depression (Darwin 1877; Woods et al. 2006). However, as was suggested by Szpiech et al. (2013), these results also imply that the damaging effect of inbreeding on fitness could be more subtle. By gathering recessive deleterious alleles in homozygous form, inbreeding allows these alleles to affect fitness. Therefore, patterns of ROH are specifically important for rare recessive alleles, because these alleles are not likely to be paired together outside of ROH. Furthermore, it suggests that recent inbreeding can shift the zygosity of deleterious alleles in only a few generations. In conservation genetics, a classic example is the Isle Royale gray wolf population, highly inbred because of isolation and small population size as well as an extreme bottleneck, and which suffers from inbreeding depression due to the presence of a high number of deleterious homozygotes in their genomes (Räikkönen et al. 2009; Hedrick et al. 2014; Robinson et al. 2019).

### ROH have proportionally less LOF homozygotes

In the second analysis, we found a significantly lower fraction of LOF homozygotes falling within short and medium ROH as compared to tolerated ones in individuals having high levels of ROH. This observation suggests that they might have been purged by natural selection. Furthermore, we do not find a significantly different fraction of LOF homozygotes in individuals with high levels of long ROH compared to tolerated ones. As these ROH are likely comprised of young haplotypes paired via inbreeding, this pattern could be the result of purifying selection not having enough time to purge these alleles. We note that it could also be due to sampling variance. Overall, these results are consistent with the idea that inbreeding can facilitate the purging of highly deleterious variants from populations over time. This has been observed in other genomic studies, including mountain gorillas (Xue et al. 2015), Indian tigers (Khan et al. 2021), Alpine ibex (Grossen et al. 2020) or other species (Ochoa and Gibbs 2021; Xie et al. 2022; Kleinman-Ruiz et al. 2022).

It is important to note that our results were obtained with a small set of LOF variants (n=179), which reduces the power of our analysis. Furthermore, the potential positive effects of inbreeding should be conditioned by several factors, such as that the population does not suffer from local extinction due to demographic stochasticity during the bottleneck, that the overall fitness is not affected too much by recessive deleterious alleles not purged effectively (Robinson et al. 2023), or that the overall lack of genetic diversity does not weaken population adaptive potential facing future environmental challenges.

### Some deleterious variants are maintained via genetic hitchhiking

When looking for signs of high-frequency deleterious variants that would be maintained by genetic hitchhiking (or, possibly, by “deleterious sweeps” (Johri et al. 2021)), we found several mutations across the five subspecies that fell within regions under selection. Because variants classified by SIFT as deleterious are mutations that are likely to affect protein function (Vaser et al. 2016), these mutations could be either advantageous or detrimental for the individual fitness. In the case of *M. m. mulatta, M. m. lasiotis, M. m. littoralis* and *M. m. tcheliensis*, each of the variants we identified is alone on its respective chromosome. In the case of *M. m. brevicaudus* however, we identified four deleterious mutations on chromosome 14 in a 1 Mb region in the vicinity of BIRC2. It may be that these variants are under positive selection, or it could be that one or more of them are detrimental and are maintained by genetic hitchhiking. We note that BIRC2 was also identified to be under positive selection by Liu et al. (2018) and could be the gene driving the other variants to high frequency.

We do not however find any more overlap with the genes found to be under selection by Liu et al. (2018). This could be due to the different filters we applied on our data, or the different methods we used to predict mutations effect and to detect selection. In particular, Liu et al. (2018) used deleterious allele predictions that were inferred from human data and then lifted over to the macaque genome, whereas we predicted variant effect on the macaque genome directly. Furthermore, to identify putatively deleterious variation that may be hitchhiking to high frequency, we used a selection scan method (nSL), which is best powered for on going (pre-fixation) positive selection, whereas Liu et al. (2018) used pairwise *F*_*ST*_ which is best powered for post-fixation sweeps (Szpiech et al. 2021).

### Implications for conservation research

Though rhesus macaques are not classified as an endangered species (Singh et al. 2020), conservation actions in other non-human primates and, more generally, endangered vertebrates can benefit from this work. To limit the effects of genetic diversity loss in small and isolated endangered populations, a common genomic-informed conservation strategy is genetic rescue, which consists in introducing novel genetic variation via gene flow, for instance by translocating animals (Whiteley et al. 2015; Tallmon et al. 2004; Hedrick and Garcia-Dorado 2016). Although there is little empirical evidence that the accumulation of deleterious mutations necessarily leads to extinction (Robinson et al. 2023), there is some evidence that total genomic ROH levels can affect fitness (Stoffel et al. 2021; Swinford et al. 2023), presumably because they are proxies for genetic load. The effects of strongly deleterious (LOF) homozygotes on extinction risks in highly inbred populations have been observed several times (Robinson et al. 2019; Johnson et al. 2010). Understanding the dynamics of deleterious and LOF mutations and therefore genetic load is crucial to implement effective genetic rescue strategies (Dussex et al. 2023; Kyriazis et al. 2021).

Effective conservation strategies should try to minimize homozygosity to mask harmful mutations, as proposed by Bossu et al. (2023) who included homozygosity information in their assisted breeding program for burrowing owls. Intentional outbreeding may even lead to heterosis (Whiteley et al. 2015). Indeed, it has been suggested that heterosis explains the beneficial outcomes of genetic rescue (Robinson et al. 2023; Kyriazis et al. 2021), though these positive effects may persist over only a few generations (Tallmon et al. 2004). In the case of historically small and isolated populations, animal translocations from larger populations should be considered carefully as it could potentially lead to the introduction of strongly deleterious (LOF) recessive variants that may manifest as homozygotes in high inbreeding contexts. A classic example of such a scenario is the Isle Royale wolf population, that dramatically collapsed after the arrival of a single migrant in 1990, who is thought to have introduced strongly recessive deleterious alleles on the island population (Hedrick et al. 2014; Robinson et al. 2016, 2019). In such cases and as emphasized by Kyriazis et al. (2021), genetic rescue strategies should focus on limiting strongly deleterious variation rather than increasing overall genetic diversity.

## Conclusion

In this study, we have analyzed the genome of 79 wild Chinese rhesus macaques to investigate the maintenance of deleterious and LOF variation in natural populations. We have shown that recent inbreeding is an important driver in the gathering of deleterious alleles in homozygous state. On the other hand, we have found signals of purging of LOF alleles in ROH, indicating that inbreeding could allow natural selection to remove recessive strongly deleterious variants from natural populations. Finally, our study suggests that some deleterious variants have also been maintained at high frequency in some populations by genetic hitchhiking. Our results add understanding to how harmful alleles are transmitted and maintained in wild non-human primates, and how evolutionary processes such as demography and natural selection can shape the distribution of harmful variation. This work is particularly relevant in the context of conservation, as there is a need to better understand how mildly and strongly deleterious mutations affect the health and viability of small and isolated endangered populations to implement effective conservation strategies such as genetic rescue.

## Material and Methods

### Data preparation

Whole genomes of 79 wild-born Chinese rhesus macaques were sequenced by Liu et al. (2018) from 17 sites in China, across the five different subspecies: *M. m. brevicaudus* (n=5), *M. m. mulatta* (n=10), *M. m. littoralis* (n=28), *M. m. lasiotis* (n=31) and *M. m. tcheliensis* (n=5). In their study, Liu et al. (2018) also included two individuals retrieved from the National Center for Biotechnology Information (NCBI) that we chose to exclude. Liu et al. (2018) called all multiallelic polymorphic sites following GATK best practices using the reference genome rheMac8, and identified 52,534,348 passing polymorphic autosomal sites. For our analysis, we filtered multiallelic SNPs and indels, and set genotypes with a genotype quality (GQ) lower than 20 to missing. We then filtered sites with more than 10% missing genotypes across the 79 individuals. An updated reference genome assembly for *M. mulatta* was recently released, increasing the sequence contiguity 120-fold and providing a better understanding of the rhesus macaque genome (Warren et al. 2020). For this reason, we decided to lift over our variants on the reference genome Mmul_10, which was achieved using LiftOver from the University of California Santa Cruz (UCSC) Genome browser.

### Annotating variants with SIFT

Variant effect was predicted using SIFT 4G (Vaser et al. 2016). SIFT (Sorting Intolerant From Tolerant) For Genomes is an algorithm that classifies a variant as synonymous or non-synonymous (missense, stop-loss, stop-gain, stop-loss), and predicts whether the amino acid substitution is deleterious to protein function based on evolutionary conservation information. No database was available for the reference genome Mmul_10 so we created it using the genome and gene annotations from UCSC Genome Browser, and the UniRef90 protein data base (Uniprot). We used the multi-transcript option, meaning that for one substitution, the effect on all transcripts were predicted. We filtered variants with the “WARNING! Low confidence” annotation, and when two or more transcripts of the same gene were given opposite predictions, we kept the worst prediction. Furthermore, SIFT does not provide a prediction for variants identified as stop-loss, stop-gain and start-loss and we therefore manually classified them as LOF, as we can expect the loss of a stop codon (stop-loss) potentially resulting in a transcript read-through error, the presence of a premature stop codon (stop-gain) potentially resulting in a truncated transcript, or the loss of the initiation codon (start-loss) potentially affecting the transcription of the transcript itself, to have a high chance of negatively affecting protein function. While each of these mutation types may not truly result in a LOF effect (e.g., nonsense mediated decay can prevent some truncated transcripts from being translated), we expect that this set of mutations as a whole generally represents mutations that are likely to be strongly deleterious (Lek et al. 2016).

### Calling ROH

ROH were called using the program GARLIC v1.1.6a, a model-based approach that was implemented by Szpiech et al. (2017). In addition to performing ROH calling, GARLIC also classifies these ROH into classes according to their length (Szpiech et al. 2017). It relies on an ROH calling algorithm introduced by Pemberton et al. (2012) that uses the logarithm of the odds (LOD) score measure of autozygosity in a sliding-window along the genome (Szpiech et al. 2017; Pemberton et al. 2012). We let GARLIC choose the window size, starting with a window size of 100 nucleotides and increasing the size by 10 nucleotides until converging, and we let GARLIC choose for the number of ROH length classes. Furthermore, we set the number of re-samples to 40 to infer allele frequencies in order to mitigate biases from unequal sample sizes. Finally, the centromere positions for built Mmul_10 were retrieved from the UCSC genome browser and provided to GARLIC in order to ensure ROH that spanned these regions were not called. All other parameters were set to default values, and we called ROH on the 79 individuals (garlic flags –auto-winsize –auto-overlap-frac –winsize 100 –tped input.tped –tfam input.tfam –tgls input.tgls –gl-type GQ –resample 40 –centromere centromere_file.txt –out output_file.txt). For each individual, we obtained the list of ROH with their position on the genome (chromosome, beginning and end), the size class they were classified into, and their length.

### Mapping variants onto ROH

We split our subset of variants between tolerated, deleterious and LOF substitutions. Then we filtered the homozygotes for the reference allele and the heterozygotes. We intersected these sites with ROH regions using BEDTools (Quinlan and Hall 2010). For each category of variants, we counted the number of homozygotes falling inside any size ROH and each size class individually, and outside of them.

### Statistical analysis of variation distribution

*G*_*i,j*_, the total fraction of the genome covered by any ROH (*j* = *R*), short (*j* = *A*), medium (*j* = *B*) or long (*j* = *C*) ROH in individual *i* (fig. 2), was computed as:

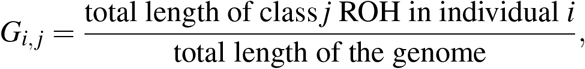

with the total length of the genome equal to 2.9 Gb (Warren et al. 2020).

The fractions of tolerated 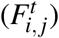, deleterious 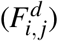 and LOF 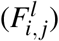 homozygotes occurring within ROH of type *j* were computed as:

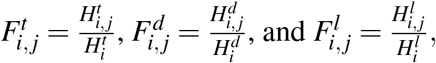

where 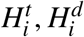, and 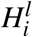 are the total number of tolerated, deleterious and LOF homozygotes genome-wide, respectively, and 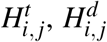, and 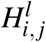 are the total number of tolerated, deleterious and LOF homozygotes, respectively, falling within ROH of type *j*, with *j* = *R* for any size ROH, *j* = *A* for short ROH, *j* = *B* for medium ROH and *j* = *C* for long ROH.

The statistical significance between the separate regressions for deleterious and tolerated homozygotes and any size ROH (fig. 4A) was assessed with the following linear model:

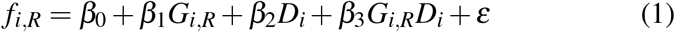

where *f*_*i,R*_ is a vector of length 158 containing, for all individuals, the fraction of genome-wide deleterious homozygotes in any ROH region 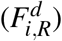 and the fraction of genome-wide tolerated homozygotes in any ROH region 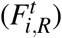. *G*_*i,R*_ is the fraction of the genome covered by any size ROH in individual *i*, and *D*_*i*_ is an indicator variable taking the value 1 if the response variable is calculated from predicted deleterious variation and 0 if the response variable is calculated from predicted to be tolerated variation. Thus, a statistically significant *β*_2_ implies a difference in the intercepts of separate regression for deleterious and tolerated homozygotes, and a statistically significant *β*_3_ indicates a significant difference in slope.

Similarly, we assessed the statistical difference between the two regressions for short, medium and long ROH (fig. 4B, C and D) with the following linear model:

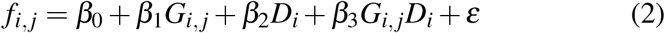

where the variables are defined as above for short (*j* = *A*), medium (*j* = *B*) and long (*j* = *C*) ROH.

### Identifying regions under selection

We identified regions under selection independently for each subspecies with the program selscan (Szpiech and Hernandez 2014). Because selscan does not handle missing data, we first phased our data using the program SHAPEIT2 v904 (Delaneau and Marchini 2014) to recover part of missing genotypes. For a lack of genetic map for Mmul_10, we assumed a recombination rate constant over the genome using the –rho option with a value of 0.00017043. This value was computed assuming a recombination rate per site per generation of 5.126 × 10^−9^ (Szpiech et al. 2021), and the average effective population size over the 5 subspecies of rhesus macaques using our estimates of genetic diversity (Supplementary Table S4) and assuming a mutation rate of 1.08 × 10^−8^ (Liu et al. 2018). We then ran selscan (Szpiech and Hernandez 2014) using the nSL statistic (Ferrer-Admetlla et al. 2014) (selscan flags –nsl –thread 8 –vcf file.vcf). Next, we normalized the scores in frequency bins using the genome-wide empirical background with selscan’s norm function (norm flags –nsl –files file.vcf –bp-win) (Szpiech and Hernandez 2014). We kept the windows with a normalized nSL score in the top 1%. Chromosomal regions under selection were then annotated with the Ensembl Genes database (Martin et al. 2023) using the UCSC Table Browser (Nassar et al. 2023; Karolchik et al. 2004).

### Identifying high frequency deleterious variants in regions under selection

Deleterious variants present in the regions under selection were identified using BEDTools (Quinlan et al. 2010). We then computed the site-frequency-spectrum (SFS) on variants inside and outside these regions. We identified the sites at high frequency by keeping the alternate alleles present at a frequency equal or higher than 0.7 in each subspecies. These sites were annotated with the Ensembl database, and the genes ontologies were extracted with PANTHER (Thomas et al. 2022). To be able to compare the SFS of the different subspecies (see fig. 4), we re-sampled the SFS with the smallest sample size, 10 haploid individuals, using the following equation:

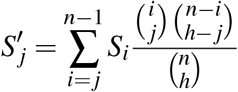

where *S* is the original SFS of size *n* and *S*_*i*_ the number of snps carried by *i* (haploid) individuals, *S*^′^ is the re-sampled SFS of size *h* and *S*^′^_*j*_ the number of variants carried by *j* individuals, and with *h* ≤ *n*.

## Data Availability

Macaque whole genome VCFs are available at http://doi.org/10.5524/100484. All code is available at https://github.com/camillesteux/rhesusmacaques.

## Acknowledgements

This work was supported by the National Institute of General Medical Sciences of the National Institutes of Health under Award Number R35GM146926 (ZAS) and by start-up funds from the Pennsylvania State University’s Department of Biology (ZAS). Computations for this research were performed using the Pennsylvania State University’s Institute for Computational Data Sciences’ Roar supercomputer.

## References

Agrawal AF, Whitlock MC. 2012. Mutation load: the fitness of individuals in populations where deleterious alleles are abundant. Annualreviewofecologyevolutionandsystematics 43:115–135.

Barton NH. 2000. Genetic hitchhiking. Philos Trans R Soc Lond B Biol Sci. 355:1553–1562.

Benazzo A, et al. 2017. Survival and divergence in a small group: The extraordinary genomic history of the endangered apennine brown bear stragglers. Proc Natl Acad Sci USA. 114:E9589– E9597.

Bertorelle G, et al. 2022. Genetic load: genomic estimates and applications in non-model animals. Nat Rev Genet. 23:492–503.

Bossu CM, et al. 2023. Genomic approaches to mitigating genetic diversity loss in declining populations. MolecularEcology 32:5228–5240.

Brüniche-Olsen A, Kellner KF, Anderson CJ, DeWoody JA. 2018. Runs of homozygosity have utility in mammalian conservation and evolutionary studies. Conserv Genet. 19:1295–1307.

Ceballos FC, Joshi PK, Clark DW, Ramsay M, Wilson JF. 2018. Runs of homozygosity: windows into population history and trait architecture. Nat Rev Genet. 19:220–234.

Charlesworth B, Jensen JD. 2021. Effects of selection at linked sites on patterns of genetic variability. Annualreviewofecologyevolutionandsystematics 52:177–197.

Charlesworth D, Willis JH. 2009. The genetics of inbreeding depression. Nat Rev Genet. 10:783– 796.

Darwin C. 1877. The effects of cross and self fertilisation in the vegetable kingdom. D. Appleton.

de Manuel M, et al. 2020. The evolutionary history of extinct and living lions. Proceedingsoft-heNationalAcademyofSciences 117:10927–10934.

Delaneau O, Marchini J. 2014. Integrating sequence and array data to create an improved 1000 genomes project haplotype reference panel. Naturecommunications 5:3934.

Dussex N, Morales HE, Grossen C, Dalén L, van Oosterhout C. 2023. Purging and accumulation of genetic load in conservation. Trendsinecologyandevolution.

Ferrer-Admetlla A, Liang M, Korneliussen T, Nielsen R. 2014. On detecting incomplete soft or hard selective sweeps using haplotype structure. Mol Biol Evol. 31:1275–1291.

Gillespie JH. 2004. Population genetics: a concise guide. JHU press.

Grossen C, Guillaume F, Keller LF, Croll D. 2020. Purging of highly deleterious mutations through severe bottlenecks in alpine ibex. Nat Commun. 11:1001.

Hedrick PW, Garcia-Dorado A. 2016. Understanding inbreeding depression, purging, and genetic rescue. Trendsinecologyandevolution 31:940–952.

Hedrick PW, Kalinowski ST. 2000. Inbreeding depression in conservation biology. Annu Rev Ecol Syst. 31:139–162.

Hedrick PW, Peterson RO, Vucetich LM, Adams JR, Vucetich JA. 2014. Genetic rescue in isle royale wolves: genetic analysis and the collapse of the population. Conserv Genet. 15:1111– 1121.

Hernandez RD, et al. 2007. Demographic histories and patterns of linkage disequilibrium in chinese and indian rhesus macaques. Science 316:240–243.

Johnson WE, et al. 2010. Genetic restoration of the florida panther. Science 329:1641–1645.

Johri P, Charlesworth B, Howell EK, Lynch M, Jensen JD. 2021. Revisiting the notion of deleterious sweeps. Genetics 219:iyab094.

Kardos M, Luikart G, Allendorf FW. 2015. Measuring individual inbreeding in the age of genomics: marker-based measures are better than pedigrees. Heredity 115:63–72.

Karolchik D, et al. 2004. The ucsc table browser data retrieval tool. Nucleic Acids Res. 32:D493– D496.

Keller MC, Visscher PM, Goddard ME. 2011. Quantification of inbreeding due to distant ancestors and its detection using dense single nucleotide polymorphism data. Genetics. 189:237–249.

Khan A, et al. 2021. Genomic evidence for inbreeding depression and purging of deleterious genetic variation in indian tigers. Proc Natl Acad Sci USA. 118:e2023018118.

Kleinman-Ruiz D, et al. 2022. Purging of deleterious burden in the endangered iberian lynx. Proc Natl Acad Sci USA. 119:e2110614119.

Kyriazis CC, Wayne RK, Lohmueller KE. 2021. Strongly deleterious mutations are a primary determinant of extinction risk due to inbreeding depression. Evolutionletters 5:33–47.

Lek M, et al. 2016. Analysis of protein-coding genetic variation in 60,706 humans. Nature 536:285–291.

Leon-Apodaca AV, et al. 2023. Genomic consequences of isolation and inbreeding in an island dingo population. bioRxiv. p. 2023.09.15.557950.

Liu Z, et al. 2018. Population genomics of wild chinese rhesus macaques reveals a dynamic demographic history and local adaptation, with implications for biomedical research. GigaScience. 7:giy106.

Lynch M, Conery J, Burger R. 1995. Mutation accumulation and the extinction of small populations. TheAmericanNaturalist 146:489–518.

Martin FJ, et al. 2023. Ensembl 2023. Nucleic Acids Res. 51:D933–D941.

McQuillan R, et al. 2008. Runs of homozygosity in european populations. Am J Hum Genet. 83:359–372.

Mooney JA, Yohannes A, Lohmueller KE. 2021. The impact of identity by descent on fitness and disease in dogs. Proc Natl Acad Sci USA. 118.

Muller HJ. 1950. Our load of mutations. Americanjournalofhumangenetics 2:111.

Nassar LR, et al. 2023. The ucsc genome browser database: 2023 update. Nucleic Acids Res. 51:D1188–D1195.

Ochoa A, Gibbs HL. 2021. Genomic signatures of inbreeding and mutation load in a threatened rattlesnake. Mol Ecol. 30:5454–5469.

Ohta T. 1973. Slightly deleterious mutant substitutions in evolution. Nature. 246:96–98.

Pemberton TJ, Szpiech ZA. 2018. Relationship between deleterious variation, genomic autozygosity, and disease risk: insights from the 1000 genomes project. Am J Hum Genet. 102:658–675.

Pemberton TJ, et al. 2012. Genomic patterns of homozygosity in worldwide human populations. Am J Hum Genet. 91:275–292.

Quinlan AR, Hall IM. 2010. Bedtools: a flexible suite of utilities for comparing genomic features. Bioinformatics. 26:841–842.

Räikkönen J, Vucetich JA, Peterson RO, Nelson MP. 2009. Congenital bone deformities and the inbred wolves (canis lupus) of isle royale. Biol Conserv. 142:1025–1031.

Robinson J, Kyriazis CC, Yuan SC, Lohmueller KE. 2023. Deleterious variation in natural populations and implications for conservation genetics. Annu Rev Anim Biosci. 11:93–114.

Robinson JA, et al. 2016. Genomic flatlining in the endangered island fox. Curr Biol. 26:1183– 1189.

Robinson JA, et al. 2019. Genomic signatures of extensive inbreeding in isle royale wolves, a population on the threshold of extinction. Sci Adv. 5:eaau0757.

Rudan I, et al. 2003. Inbreeding and risk of late onset complex disease. J Med Genet. 40:925–932.

Sams AJ, Boyko AR. 2019. Fine-scale resolution of runs of homozygosity reveal patterns of inbreeding and substantial overlap with recessive disease genotypes in domestic dogs. G3: Genes Genomes Genet. 9:117–123.

Singh M, Kumar A, Kumara H. 2020. Macaca mulatta. The IUCN Red List of Threatened Species Accessed on 12 March 2024.

Spielman D, Brook BW, Frankham R. 2004. Most species are not driven to extinction before genetic factors impact them. Proc Natl Acad Sci USA. 101:15261–15264.

Stoffel MA, Johnston SE, Pilkington JG, Pemberton JM. 2021. Mutation load decreases with haplotype age in wild soay sheep. Evolutionletters 5:187–195.

Swinford NA, et al. 2023. Increased homozygosity due to endogamy results in fitness consequences in a human population. ProceedingsoftheNationalAcademyofSciences 120:e2309552120.

Szpiech ZA, Blant A, Pemberton TJ. 2017. Garlic: genomic autozygosity regions likelihood-based inference and classification. Bioinformatics. 33:2059–2062.

Szpiech ZA, Hernandez RD. 2014. selscan: an efficient multithreaded program to perform ehh-based scans for positive selection. Mol Biol Evol. 31:2824–2827.

Szpiech ZA, Novak TE, Bailey NP, Stevison LS. 2021. Application of a novel haplotype-based scan for local adaptation to study high-altitude adaptation in rhesus macaques. Evol lett. 5:408– 421.

Szpiech ZA, et al. 2013. Long runs of homozygosity are enriched for deleterious variation. Am J Hum Genet. 93:90–102.

Szpiech ZA, et al. 2019. Ancestry-dependent enrichment of deleterious homozygotes in runs of homozygosity. Am J Hum Genet. 105:747–762.

Tallmon DA, Luikart G, Waples RS. 2004. The alluring simplicity and complex reality of genetic rescue. Trendsinecologyandevolution 19:489–496.

Thomas PD, et al. 2022. Panther: Making genome-scale phylogenetics accessible to all. Protein Sci. 31:8–22.

Uniprot. 2021. Uniprot: the universal protein knowledgebase in 2021. Nucleic Acids Res. 49:D480–D489.

Van Hooft P, Getz WM, Greyling BJ, Zwaan B, Bastos AD. 2021. A continent-wide high genetic load in african buffalo revealed by clines in the frequency of deleterious alleles, genetic hitchhiking and linkage disequilibrium. Plosone 16:e0259685.

Vaser R, Adusumalli S, Leng SN, Sikic M, Ng PC. 2016. Sift missense predictions for genomes. Nat Protoc. 11:1–9.

Von Seth J, et al. 2021. Genomic insights into the conservation status of the world’s last remaining sumatran rhinoceros populations. NatureCommunications 12:2393.

Warren WC, et al. 2020. Sequence diversity analyses of an improved rhesus macaque genome enhance its biomedical utility. Science. 370:eabc6617.

Whiteley AR, Fitzpatrick SW, Funk WC, Tallmon DA. 2015. Genetic rescue to the rescue. Trendsinecologyandevolution 30:42–49.

Woods CG, et al. 2006. Quantification of homozygosity in consanguineous individuals with auto-somal recessive disease. Am J Hum Genet. 78:889–896.

Xie HX, et al. 2022. Ancient demographics determine the effectiveness of genetic purging in endangered lizards. Mol Biol Evol. 39:msab359.

Xue Y, et al. 2015. Mountain gorilla genomes reveal the impact of long-term population decline and inbreeding. Science. 348:242–245.

Zhang Q, Guldbrandtsen B, Bosse M, Lund MS, Sahana G. 2015. Runs of homozygosity and distribution of functional variants in the cattle genome. BMC Genom. 16:1–16.

