## Supplementary material for "The Maintenance of Deleterious Variation in Wild Chinese Rhesus Macaques": Figures S1-S4

### Supplementary Figures

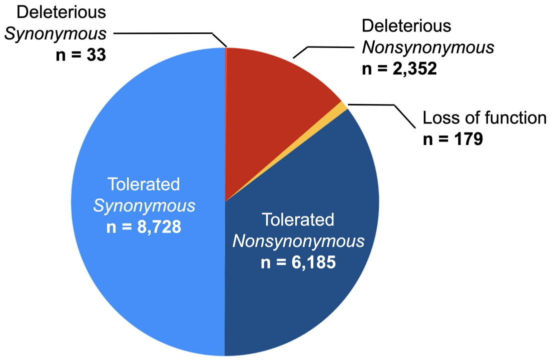


Figure S1: SIFT (Vaser et al. 2016) classification of our set of 5,713,999 after filtering. The loss-of-function category pools the variants predicted to be stop-loss, stop-gain and start-loss.

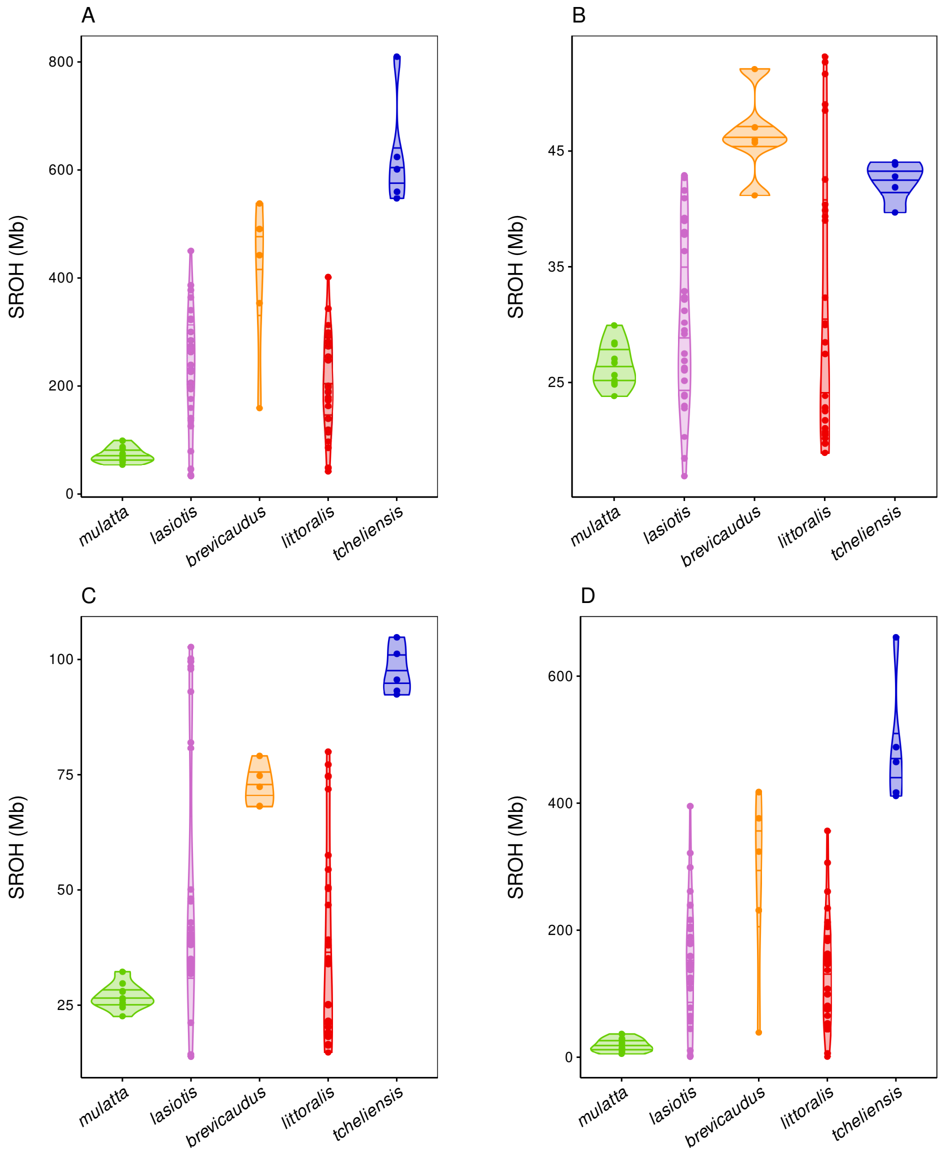


Figure S2: Distribution of the total length of ROH (SROH) identified with GARLIC (Szpiech et al. 2017) for each subspecies. (A) Any size ROH (B) short ROH (C) medium ROH and (D) long ROH.

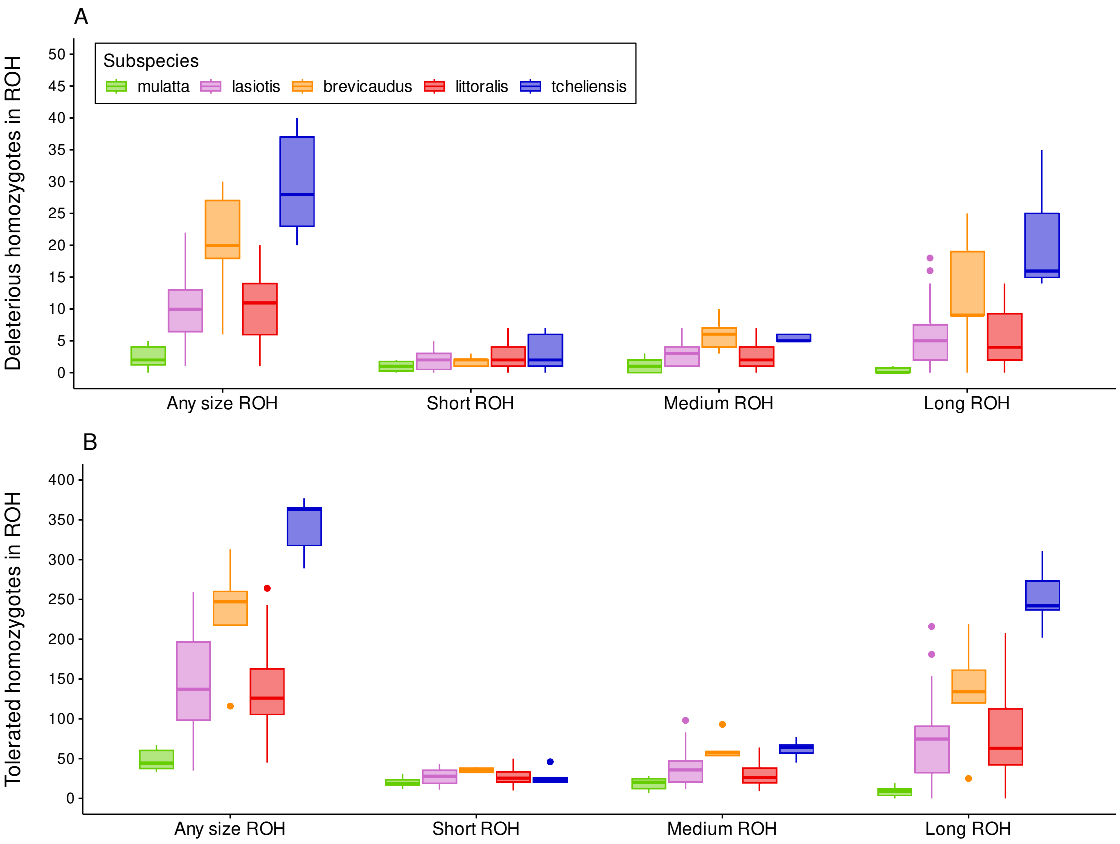


Figure S3: Number of (A) Alternate-deleterious homozygotes and (B) Tolerated-alternate homozygotes falling in any size ROH, short, medium and long ROH.

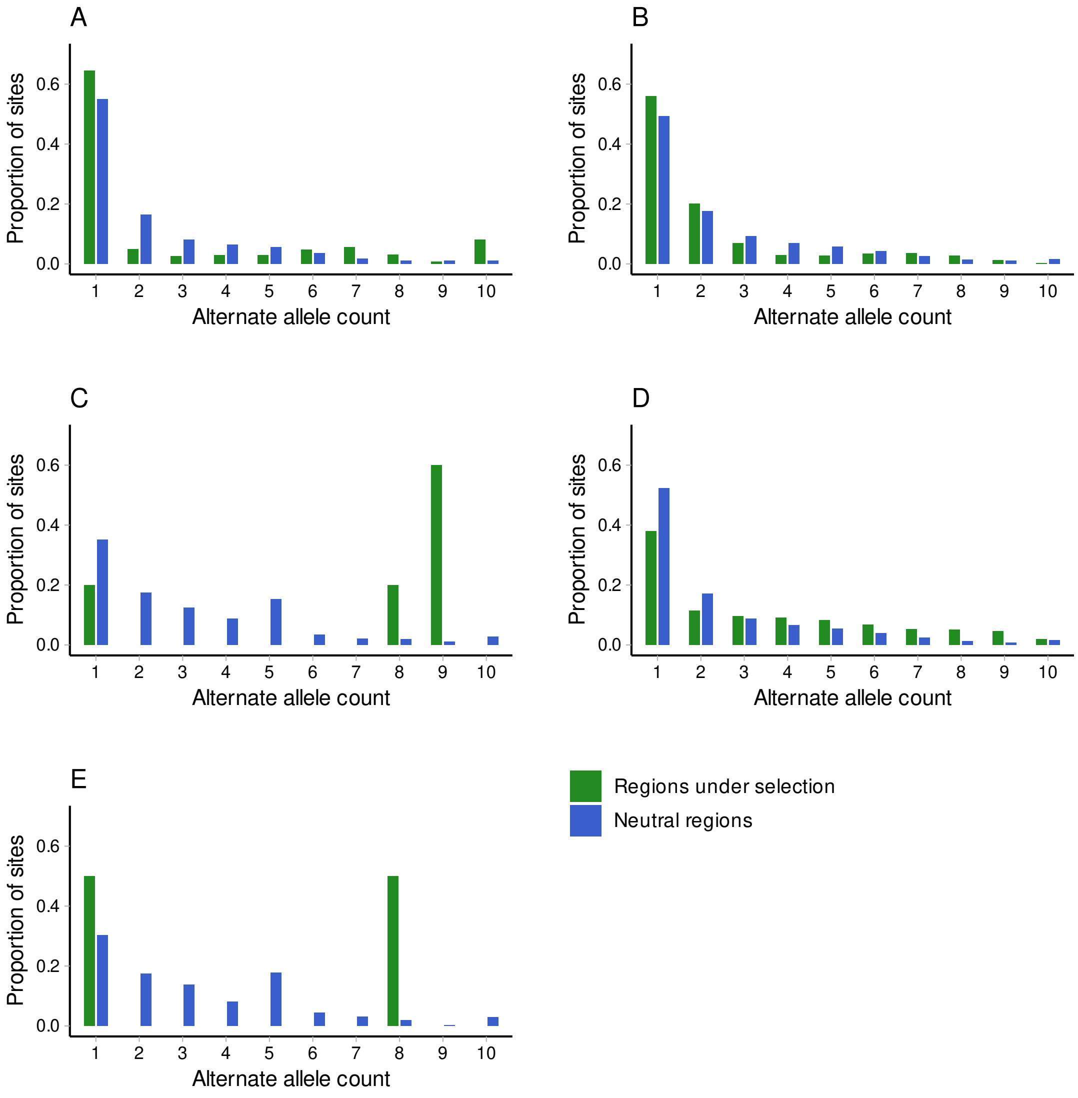


Figure S4: Site Frequency Spectrum (SFS) of deleterious variants falling in the regions under selection (green) and outside of these regions (blue) for each subspecies. SFS of *M. m. mulatta*, *M. m. lasiotis* and *M. m. littoralis* were re-sampled (n=10 haploid individuals) using the formula available in Material and Methods. (A) *M. m. mulatta* (B) *M. m. lasiotis* (C) *M. m. brevicaudus* (D) *M. m. littoralis* (E) *M. m. tcheliensis*.
